# General visual and contingent thermal cues interact to elicit attraction in female *Aedes aegypti* mosquitoes

**DOI:** 10.1101/510594

**Authors:** Molly Z Liu, Leslie B. Vosshall

## Abstract

**ABSTRACT:** Female *Aedes aegypti* mosquitoes use multiple sensory modalities to hunt human hosts to obtain a blood-meal for egg production. Attractive cues include carbon dioxide (CO_2_), a major component of exhaled breath [1, 2]; heat elevated above ambient temperature, signifying warm-blooded skin [3, 4]; and dark visual contrast [5, 6], proposed to bridge long-range olfactory and short-range thermal cues [7]. Any of these sensory cues in isolation is an incomplete signal of a human host, and so a mosquito must integrate multi-modal sensory information before committing to approaching and biting a person [8]. Here, we study the interaction of visual cues, heat, and CO_2_ to investigate the contributions of human-associated stimuli to host-seeking decisions. We show that tethered flying mosquitoes strongly orient toward dark visual contrast regardless of CO_2_ stimulation and internal host-seeking status. This suggests that attraction to visual contrast is general, and not contingent on other host cues. In free-flight experiments with CO_2_, adding a dark contrasting visual cue to a warmed surface enhanced host-seeking. Moderate warmth became more attractive to mosquitoes, and mosquitoes aggregated on the cue at all non-noxious temperatures. *Gr3* mutants, unable to detect CO_2_, were lured to the visual cue at ambient temperatures, but fled and did not return when the surface was warmed to host-like temperatures. This suggests that attraction to thermal cues is contingent on the presence of the additional human sensory cue CO_2_. Our results illustrate that mosquitoes integrate general attractive visual stimuli with the context-dependent thermal stimuli to seek promising sites for blood-feeding.

*GRAPHICAL ABSTRACT:* 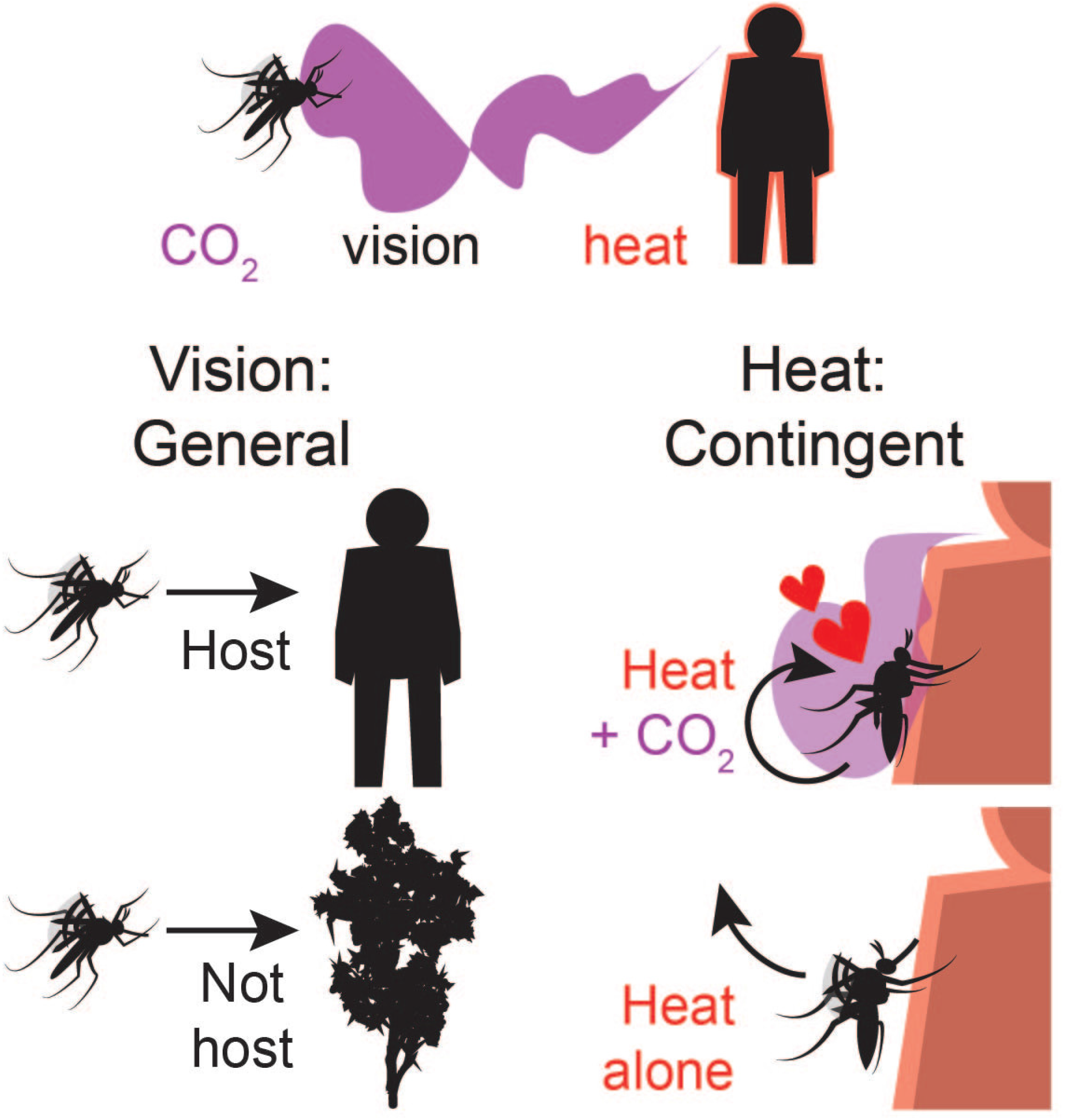

## RESULTS AND DISCUSSION

### Tethered Mosquito Visual Responses Are Not Altered by Host-Seeking State

We built a magnetic tether (“magnotether”) to study object orientation responses in mosquitoes with precise temporal and spatial control (Fig. 1A) [5, 9, 10]. The free rotation of the magnotether allows animals to control their own flight turns naturalistically, avoiding the confounds that arise from rigid-tether closed-loop experiments where the gains of visual feedback are arbitrarily chosen [11]. Because animals must be flying to be assayed in the magnotether, we focused solely on orientation in flight. In each trial, we showed an individually tethered animal a shape, such as a long dark stripe (Fig. 1B), recorded the animal’s orientation over time (Fig. 1C), computed the offset between the stimulus and animal position (Fig. 1D), and calculated the proportion of time spent with an offset within ±45° to obtain “fixation” (expected ~1.00 for attractive shapes, Fig. 1E). As controls, we presented animals with trials where only a static background was shown (“blank”) and computed offset and fixation relative to a randomly assigned fictive stimulus position (expected ~0.25, Fig. 1F-I).

**Figure 1.**
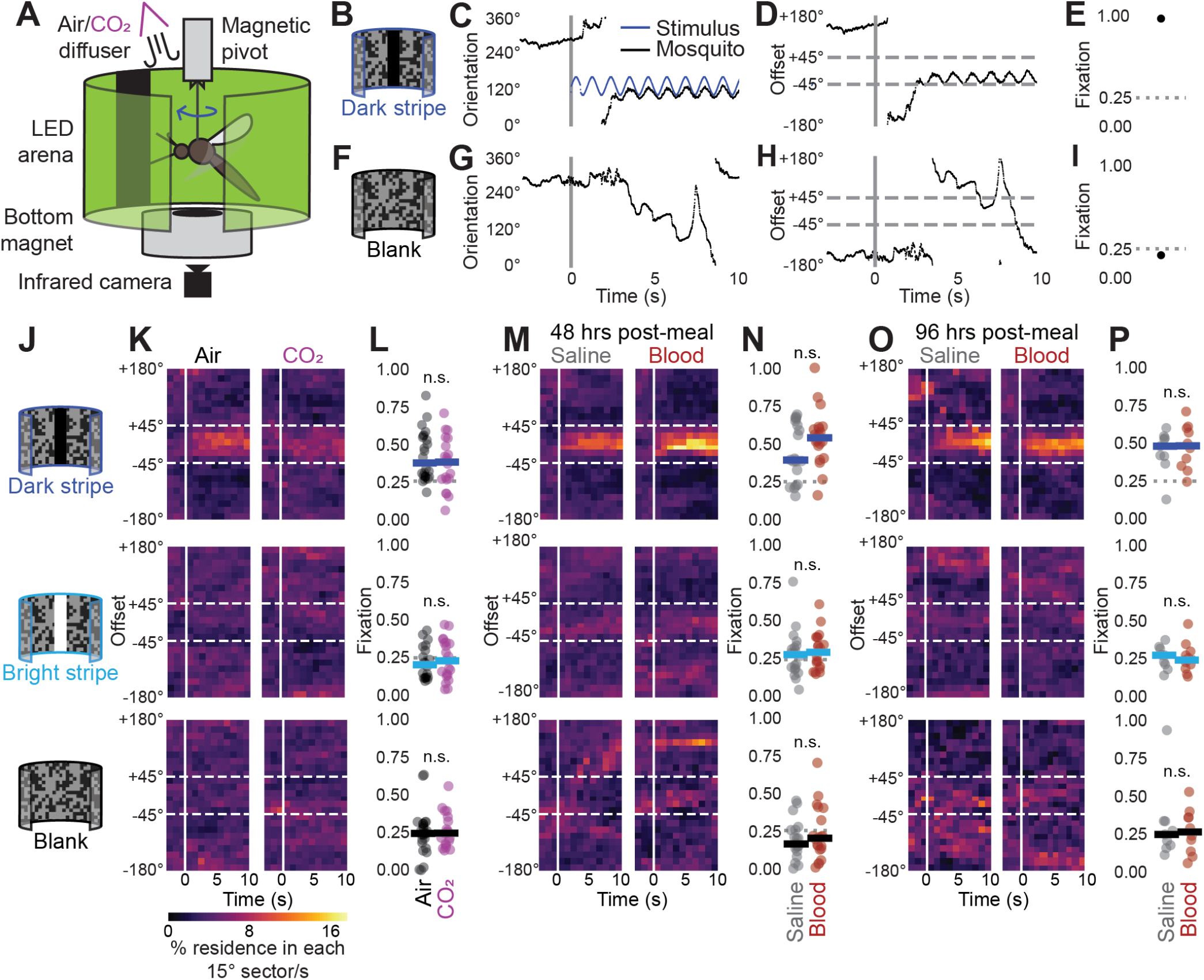
Mosquito Visual Responses are not Influenced by Host-Seeking Status. (A) Magnotether schematic. (B-I) Example magnotether traces from dark stripe (B-E) or blank (F-I) trials. (J) Magnotether stimuli. (K, M, O) Heat maps of offset, parsed by shape of stimulus, from mosquitoes tested in air (n = 24 dark and blank, n = 20 bright) or CO_2_ (n = 20 dark and blank, n = 21 bright) (K); 48 hr post-meal, saline (n = 20) and blood (n = 20) (M); and 96 hr post-meal, saline (n = 9) and blood (n = 10) (O). Each trial corresponds to a single female. (L, N, P) Fixation, parsed by shape. n.s.: not significant (p > 0.05), Mann-Whitney U test with Bonferroni correction.

To confirm that our magnotether accurately captured object responses, we presented female *Drosophila melanogaster* flies with simple dark shapes that had known response profiles in rigid-tether and free flight (Fig. S1A). Replicating previous results [12], flies oriented strongly toward the long stripe, showed weak responses to the medium stripe, and avoided the small square (Fig. S1B-C). We presented the same simple shapes to flying female *Aedes aegypti*. However, instead of the uniform background used for flies, mosquitoes saw shapes on a patterned low-light background that stabilized their flight by enabling them to receive visual feedback in turning, and that also allowed us to present both dark and bright shapes in the same environment. Consistent with previous reports [5], mosquitoes paid attention to contrast. A medium stripe was attractive only when dark and not when bright (Fig. S1D, E). Mosquitoes did not respond to the square whether it was dark or bright (Fig. S1F, G).

Having identified visual cues that elicit responses in the magnotether, we investigated if these visual responses were altered by elevated CO_2_ (Fig. 1J). Both non-directionally [8] and as a plume [1], CO_2_ is a potent signature of host breath that amplifies responses to other host cues. In the magnotether, we applied non-directional pulses of CO_2_, previously shown to increase mosquito responses to human odor and heat [8, 4]. However, we could not detect changes in visual responses of our tethered mosquitoes (Fig. 1K, L). We speculate that the flying behavior required in the magnotether represents a state of behavioral arousal that cannot be further increased by CO_2_.

Attraction to host cues is also modulated by internal state. When a female mosquito ingests a blood-meal, her previously strong attraction to host cues is suppressed until she lays eggs [13]. Conversely, mosquitoes fed saline continue to be attracted to hosts. We compared visual orientation behaviors of mosquitoes after ingestion of blood or saline, and were not able to detect a difference in visual orientation (Fig. 1M-P). Because flying mosquitoes show general, non-contingent attraction to visual cues, we suggest that dark visual contrast signals potential landing sites, as has been proposed for *Drosophila* [14].

### Host-Seeking Thermotaxis can be Enhanced by Co-Presentation of a Visual Cue

Given that responses to visual cues in the magnotether are not modulated by CO_2_ or blood-feeding, we asked if the reciprocal might be true: do visual cues affect mosquito responses to a signature host cue, such as human skin temperature? To test this, we adapted a previously described free flight thermotaxis assay [4] that measures mosquito occupancy on a temperature-controlled Peltier in the presence or absence of CO_2_. We modified the Peltier by affixing to its center a 2 cm black circle (“dot”), a visual stimulus previously shown to attract host-seeking mosquitoes in free flight [7] (Fig. 2A). We tracked the two-dimensional location of mosquitoes while setting the internal temperature of the Peltier to various temperatures (Fig. 2B). The dot did not present a detectably different thermal signature from the rest of the Peltier (Fig. 2C), meaning that any differences in occupancy should be due to visual features of the dot. We presented a heat ramp to groups of 45-50 female mosquitoes, co-presented with pulses of CO_2_ and either the dot or a blank control (Fig. 2D).

**Figure 2.**
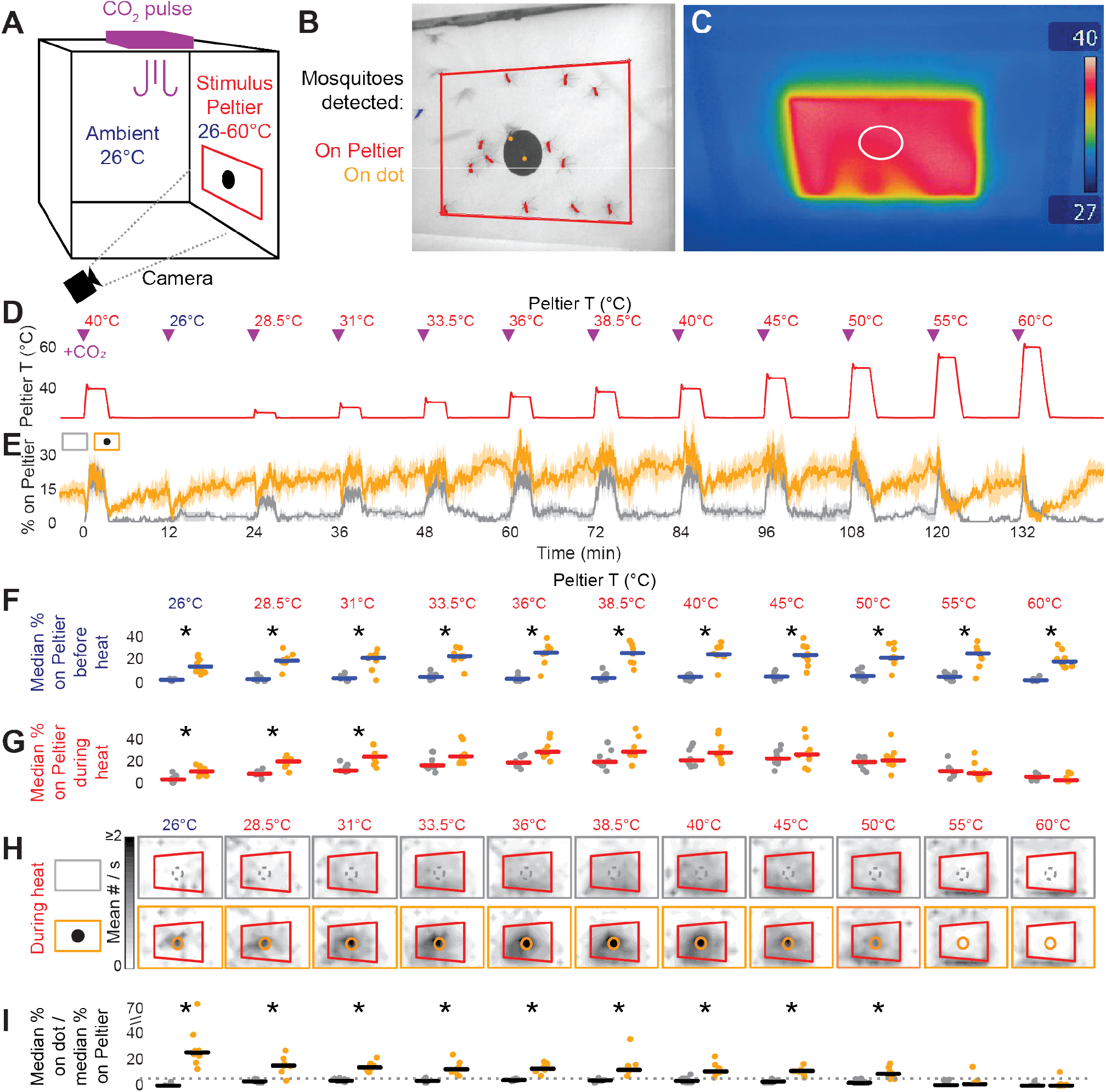
Visual Cue Enhances Mosquito Thermotaxis to Moderate Heat. (A) Schematic of heat-seeking assay with 2 cm diameter black dot as visual cue. (B) Camera image, showing mosquitoes on Peltier (red outline) sampled at 1 Hz. (C) Thermal image of the Peltier set to 40°C. White outline shows boundary of black dot. (D) Peltier temperature. CO_2_ was pulsed in for 20 s at onset of each heat bout (purple triangle). (E) Median % of mosquitoes on Peltier over entire experiment ± median absolute deviation, blank (gray, n = 8) or visual cue (orange, n = 8). (F, G, I) Median % of mosquitoes on the Peltier 90-0 s before onset of heat (“before heat”) (F), 90-180 s after onset of heat (“during heat”) (G), and on the area of the dot during heat (I), normalized by median % of mosquitoes anywhere on the Peltier (“on dot”). Dotted line in (I) indicates expected value from a uniform spatial distribution. * p < 0.05, Mann-Whitney U test with Bonferroni correction. (H) Heat maps showing mean mosquito occupancy on the Peltier (red outline) and surrounding area with blank (gray) or visual cue (orange) during seconds 90–180 of heat.

Consistent with a previous study [4], mosquito occupancy on the Peltier increased as the temperature increased to 28.5-50°C, then decreased as it heated to 55-60°C (Fig. 2E). Because a mosquito landing on the Peltier surface experiences a temperature several degrees lower than the set internal temperature, we speculate that the attractive temperatures found correspond to the temperature of human skin, approximately 29-35°C [15].

We saw two striking differences when the dot was present. First, mosquitoes occupied the Peltier at high rates at ambient temperature (26°C), including between heat bouts and before pulses of CO_2_ (Fig. 2F). This is consistent with our finding that tethered mosquitoes are attracted to dark contrast independent of elevated CO_2_ (Fig. 1, Fig. S1), as well as previously published findings showing that freely flying mosquitoes approached small black dots [7]. Second, the dot enhanced total levels of mosquito occupancy on the Peltier at warm temperatures cooler than human skin (28.5-31°C, Fig. 2G). To examine the effect of the dot more closely, we compared mosquito locations with and without the visual cue during thermotaxis (Fig. 2H). With the blank Peltier, mosquitoes spread themselves evenly across the warm Peltier, with a slight bias towards the bottom edge. However, when the dot was present, mosquitoes aggregated on the dot at all non-noxious temperatures (26-50°C, Fig. 2I). The same results were obtained when we randomized the order of the Peltier temperatures, showing that these effects do not depend on past history of thermal presentation (Fig. S2). Visual contrast thus has the effect of enhancing host-seeking, both by making moderate warmth more attractive to mosquitoes and by promoting aggregation on areas of high contrast at all attractive temperatures.

### Occupancy on Host-Like Temperatures is Contingent on Concurrent CO_2_ Exposure

Finally, the general attractiveness of the visual cue allowed us to investigate the role of CO_2_ in host-seeking. Previous reports showed that mosquito thermal attraction is enhanced by CO_2_ [3, 8, 16], which could be through at least two non-mutually exclusive mechanisms. First, CO_2_ could increase the propensity of mosquitoes to take off and remain in flight, which would increase their probability of coming close enough to a surface to sense its heat. Second, CO_2_ could alter the thermal preferences of mosquitoes so that they become attracted to temperatures they would ordinarily ignore or avoid. To test these possibilities, we used the dot to lure mosquitoes to the unheated Peltier (Fig. 2F). If mosquitoes are always attracted to host heat, then they should continue occupying a marked surface upon heating regardless of CO_2_ sensation. Conversely, if we only see thermal occupancy coincident with sensed CO_2_ elevation, then CO_2_ must be shifting mosquito thermal preferences.

CO_2_ is a pervasive and naturally-occurring stimulus that is difficult to remove from behavioral assays. In light of this, we tested responses to a heat ramp (Fig. 3A) with a dot-marked Peltier using *Gr3^ECFP/ECFP^* mosquitoes, which lack a functional CO_2_ receptor and cannot detect CO_2_ [8]. Regardless of their ability to sense CO_2_, mosquitoes accumulated on the Peltier between heat bouts (Fig. 3B), replicating our observation that CO_2_ does not alter mosquito general visual preferences in the magnotether assay (Fig. 1K, L).

**Figure 3.**
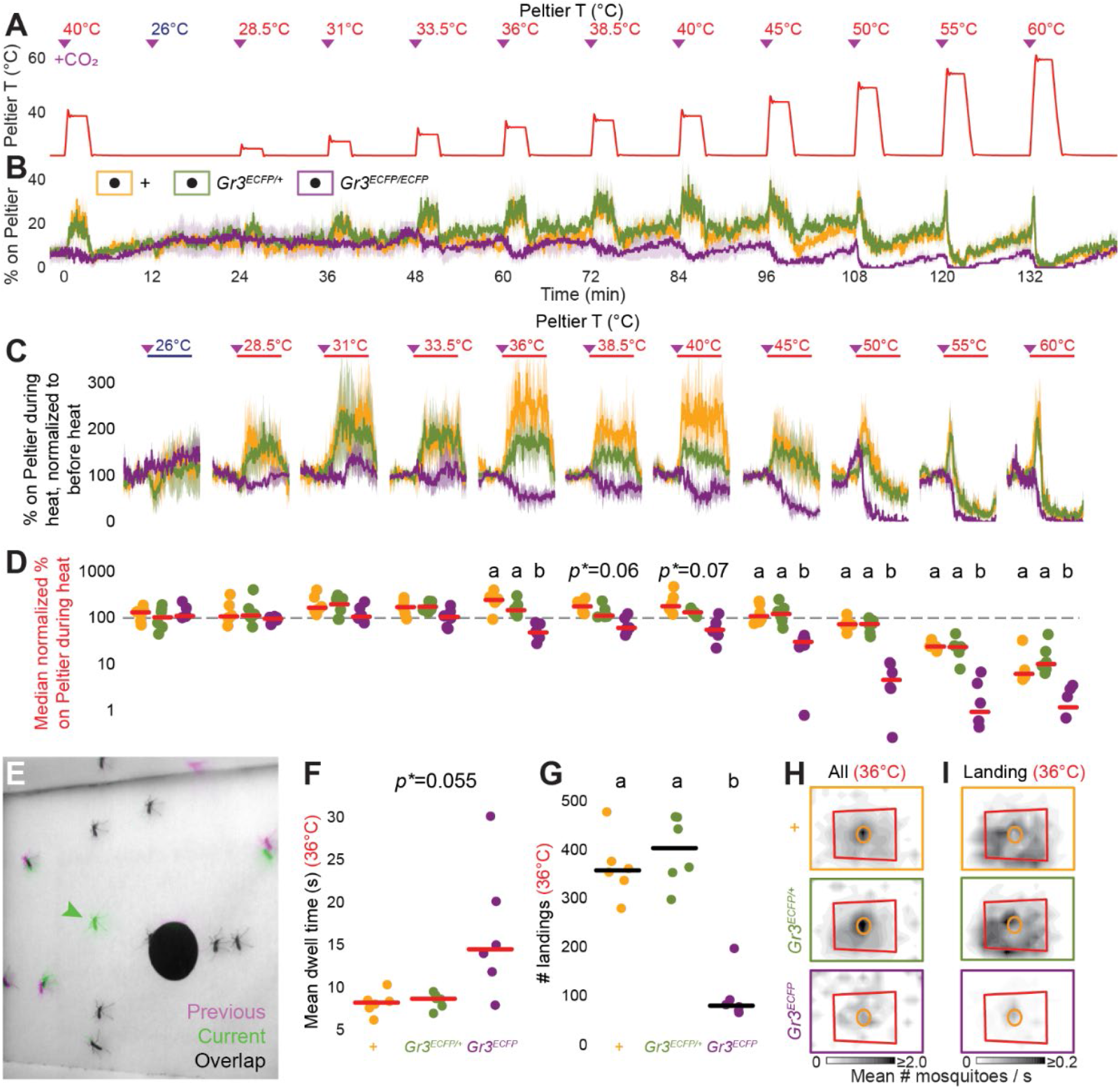
Mosquito Attraction to Host-Like Temperatures, but not to a Visual Cue, Requires CO_2_ Sensation. (A) Peltier temperature. CO_2_ was pulsed for 20 s at onset of each heat bout (purple triangle). (B) Median ± median absolute deviation % of mosquitoes on Peltier marked with a visual cue. Genotypes: *Gr3*^+/+^ (orange, n = 6), *Gr3*^*ECFP*/+^ (green, n = 6), and *Gr3^ECFP/ECFP^* (purple, n = 6). (C) Median ± median absolute deviation of % residence on Peltier over time, normalized to mean residence 60-0 s before heat. Purple triangle: CO_2_ pulse; red horizontal line: heat on. (D) Median % of mosquitoes on the Peltier during heat, normalized as in (C) and plotted on a log10 scale to highlight deviation from 100% (dotted line). (E) Example superimposed consecutive frames. Green arrowhead: landing event. (F-G) Mean dwell time (F) and number of landings (G) on Peltier wall during 36°C heat (H-I) Heat maps showing mean mosquito occupancy (H) or landing events (I) on the Peltier (red outline) and surrounding area, during seconds 90–180 of 36°C heat. Visual cue is marked in orange. In D-I, each trial involved 45-50 female mosquitoes. In D, F, G significance (p < 0.05) was assessed with Kruskal-Wallis test with Bonferroni correction, then post hoc Mann-Whitney U test with Bonferroni correction. Data marked with different letters are significantly different from each other.

However, whereas wild-type and *Gr3^ECFP/+^* heterozygous mosquitoes increased Peltier occupancy with increasing temperatures, *Gr3^ECFP/ECFP^* mutant mosquitoes left the Peltier when it was heated to host-like temperatures (Fig. 3C, D). We observed the same decrease in occupancy in wild-type mosquitoes tested without CO_2_ pulses (Fig. S3), indicating that this effect is likely a general consequence of CO_2_ perception. We speculate that CO_2_ shifts mosquito thermal preferences in a manner independent of visual stimuli, causing mosquitoes to pursue elevated heat only when they have corroborating evidence that the heat comes from a breathing host.

CO_2_ could act to increase occupancy on host-like temperatures by causing mosquitoes to dwell on heat longer, or to land more frequently on heat. To examine how these rates changed, we manually tracked landings on and take-offs from the Peltier set to 36°C. This temperature in combination with CO_2_ attracts wild-type and *Gr3^ECFP/+^* heterozygous mosquitoes but repels *Gr3^ECFP/ECFP^* mosquitoes (Fig. 3E), allowing us to compute a population mean dwell time on heat using times of landings and take-offs.

Interestingly, wild-type and *Gr3*^*ECFP*/+^ heterozygous mosquitoes did not dwell longer on heat than *Gr3^ECFP/ECFP^* mosquitoes (Fig. 3F), perhaps because CO_2_-insensitive mosquitoes do not increase their general motor activity after CO_2_ pulses. This suggests that 36°C heat alone is an aversive cue for a landed mosquito, replicating previous findings in which mosquitoes walking on a thermal gradient avoided temperatures above 30°C [4, 17]. Instead, the increase in occupancy primarily comes from increased landing of CO_2_-sensitive mosquitoes on host-like temperatures (Fig. 3G). Furthermore, in CO_2_-sensitive mosquitoes the number of landings was five to ten times greater than the number of mosquitoes, indicating that individual mosquitoes must be landing and taking off multiple times. *Gr3^ECFP/ECFP^* mosquitoes also still landed on 36°C, but this may be visually driven. While CO_2_-sensitive mosquitoes landed evenly across the heated surface, *Gr3^ECFP/ECFP^* mosquitoes landed overwhelmingly on or around the dot (Fig. 3H, I). Mosquito attraction to host heat is thus contingent on the perception of elevated CO_2_, which drives mosquitoes to repeatedly return to the thermal stimulus.

## CONCLUSIONS

In this study, we show that both visual contrast and warm heat contribute to mosquito attraction during host-seeking. However, whereas visual contrast is generally attractive even outside the context of host-seeking, attraction to heat is contingent upon co-presentation of the chemosensory host-cue CO_2_. We speculate that this attraction to visual contrast in flight may be because flying is an energetically expensive activity [18], so finding a dark landing location may provide general respite and camouflage. However, residence on high heat brings with it threats of damage and desiccation. Indeed, mosquitoes die after 30 min exposures to air temperatures above 42°C [19]. Mosquitoes may balance this thermal threat of heat against the beneficial use of heat as a potent signature of warm-blooded hosts by only thermotaxing when they detect other reliable host cues such as CO_2_.

All animals must dynamically weigh their sensory landscape to make decisions. Studies of *Drosophila* species have described the serial sensory modules that underlie behaviors from courtship [20, 21, 22] to flight navigation [23, 24]. Here we show that *Aedes aegypti* mosquitoes use serial sensory modules in host-seeking: They fly toward visual contrast, and then sense CO_2_ to unlock thermotaxis towards potential hosts. Mosquitoes across the *Culicidae* family display an impressive variety of host choices, from mammals to cold-blooded frogs [25] to annelids [26], and the algorithms they use to weigh sensory host cues likely vary just as much. Our results illustrate how such weighting is performed in one species, providing a first glimpse into how general and contingent cues are integrated to produce host-seeking behavior in mosquitoes. With the rapid development of genetic and neuroscience tools in mosquitoes [27-30], we are poised to uncover the neuronal mechanisms underlying multimodal integration in these charismatic and deadly insects.

## SUPPLEMENTAL INFORMATION

**Figure S1.**
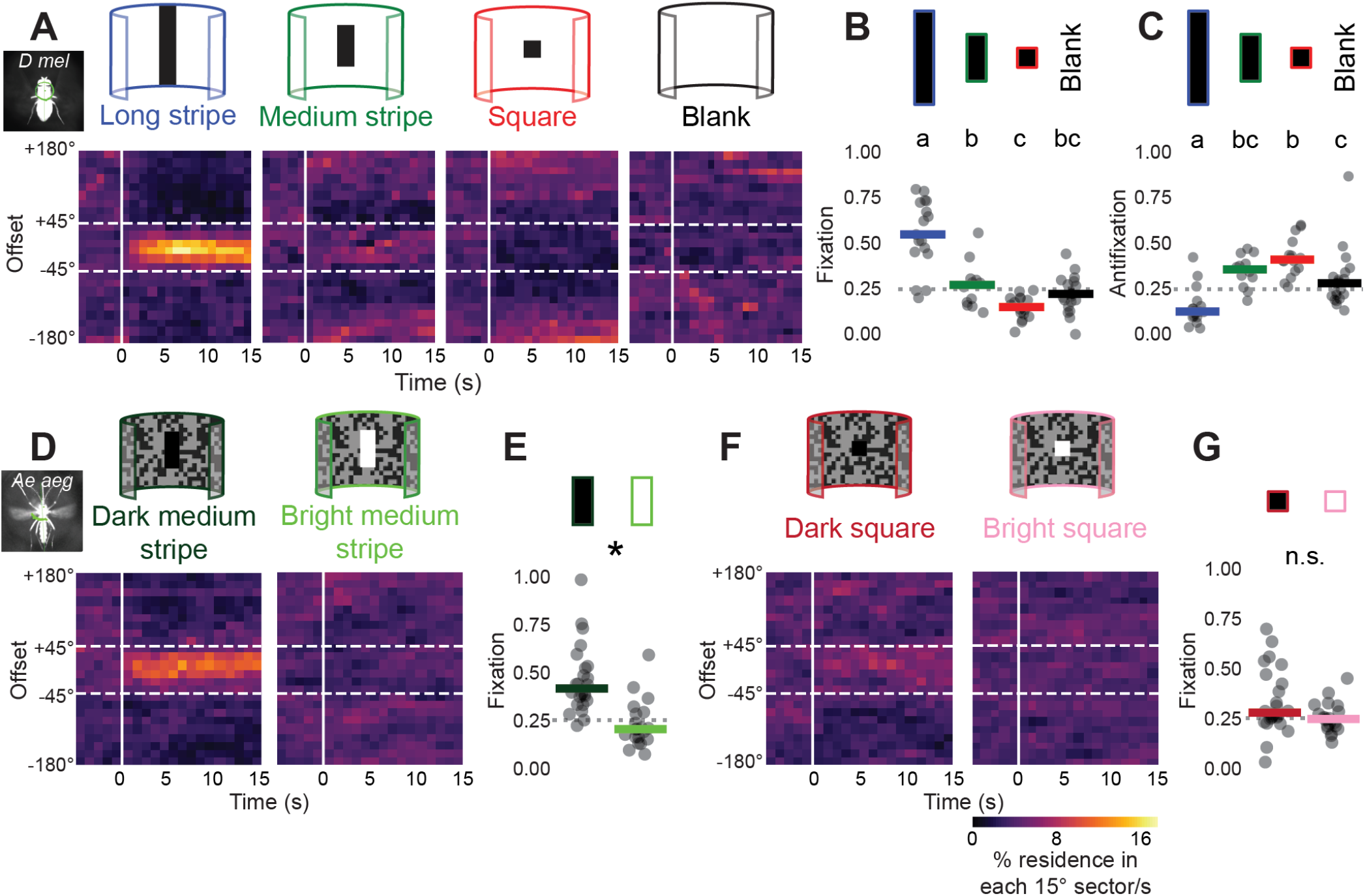
Visual Preferences of Magnotethered Vinegar Flies and Mosquitoes. (A,D,F) Heat maps of offset, parsed by shape of stimulus, from female *Drosophila melanogaster*, long stripe (n = 17), medium stripe (n = 12), square (n = 16), and blank (n = 21) (A) or female *Aedes aegypti*, dark (n = 24) and bright (n = 21) (D,F). (B,E,G) Fixation, parsed by shape. Data labeled with different letters are different from each other (p < 0. 05), Kruskal-Wallis test and post hoc Mann-Whitney U test with Bonferroni correction (B) or Mann-Whitney U test with Bonferroni correction (E,G). (C) Anti-fixation, calculated similarly to fixation but with an offset window >±135°. In A-G, each trial corresponds to a single female.

**Figure S2.**
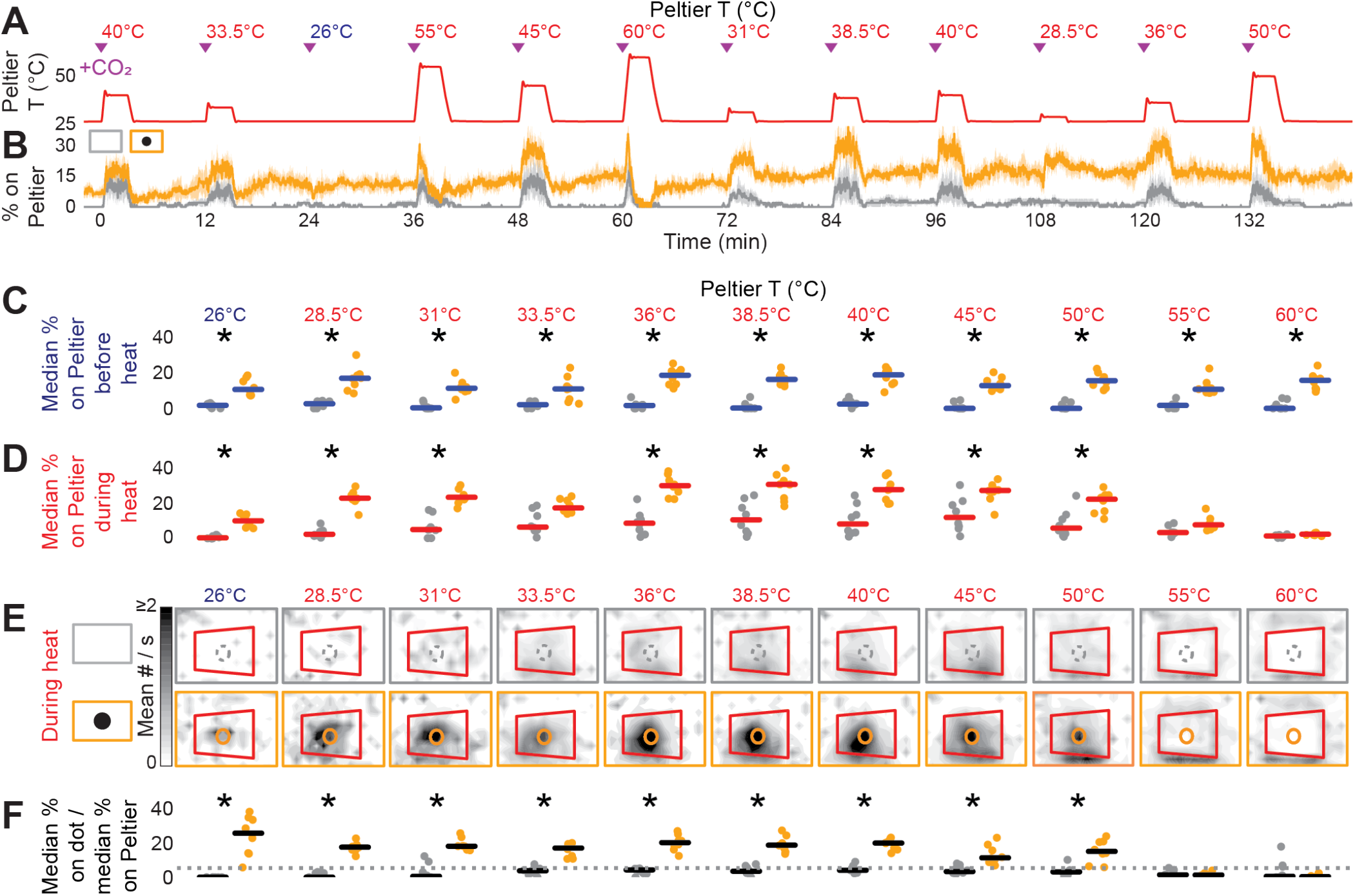
Visual Cue Enhances Mosquito Thermotaxis to Moderate Heat Presented in Shuffled Order and to Areas of Contrast. (A) Peltier temperature. CO_2_ was pulsed in for 20 s at onset of each heat bout (purple triangle). (B) Median ± median absolute deviation % of mosquitoes on Peltier, blank (gray, n = 8) or marked with circle (orange, n = 8). (C,D,F) Median % of mosquitoes on the Peltier 90-0 s before onset of heat (C), 90-180 s after onset of heat (D), and on the dot 90-180 s after onset of heat, normalized by median % of mosquitoes anywhere on the Peltier (F). Dotted line in (F) indicates expected value from a uniform spatial distribution. *p < 0.05, Mann-Whitney U test with Bonferroni correction. (E) Smoothed histograms of mosquito positions during heat, boundaries of Peltier (red) and blank (gray) or visual cue (orange). In B-F, each trial involved 45-50 female mosquitoes.

**Figure S3.**
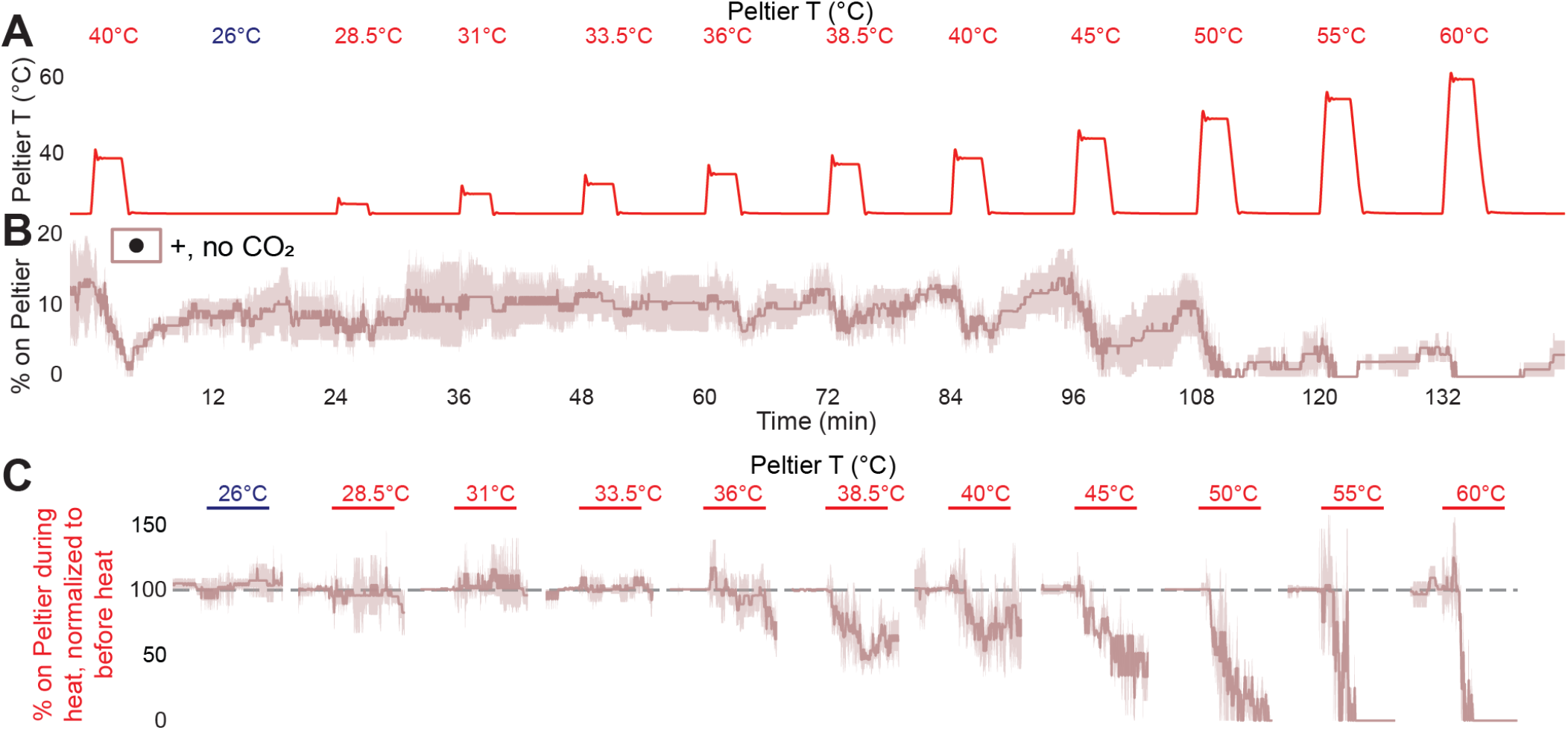
Wild-type mosquitoes are not attracted to host heat without elevated CO_2_. (A) Peltier temperature. (B) Median ± median absolute deviation % of wild-type mosquitoes (n = 6, 45-50 mosquitoes/trial) on the visually marked Peltier. (C) Median ± median absolute deviation of % residence on Peltier over time, normalized to mean residence 60-0 s before heat. Red horizontal line: heat on.

## ACKNOWLEDGMENTS

We thank Román Corfas, Caroline Kittredge Faustine, Lisa Fenk, Kaela Mei-Shing Garvin, Itzel Ishida, Cheng Lyu, Gaby Maimon, Vanessa Ruta, Cameron Toy, Waring Trible, and members of the Vosshall lab for discussion and comments on the manuscript; Lisa Fenk, Jonathan Green, Jonathan Hirokawa, and Gaby Maimon for magnotether discussions, equipment, and technical advice; Takeshi Morita for technical advice on the heat-seeking experiments; and Gloria Gordon for expert mosquito rearing. L.B.V. is an investigator of the Howard Hughes Medical Institute.

## AUTHOR CONTRIBUTIONS

Supervision and Funding Acquisition, L.B.V; Investigation, M.Z.L.; Conceptualization, Methodology, Writing, M.Z.L. and L.B.V.

## DECLARATION OF INTERESTS

The authors declare no competing interests.

## MATERIALS AND METHODS

### EXPERIMENTAL MODEL AND SUBJECT DETAILS

#### Fly rearing and maintenance

*Drosophila melanogaster* wild-type laboratory strains (isoD) were maintained and reared with 25°C, with a photoperiod of 12 hr light:12 hr dark on standard corn-meal agar. Flies were sexed and sorted under cold anesthesia (4°C).

#### Mosquito rearing and maintenance

*Aedes aegypti* wild-type laboratory strains (Orlando) and mutant strains were maintained and reared at 25-28°C, 70%–80% relative humidity with a photoperiod of 14 hr light:10 hr dark (lights on at 7 a.m.) as previously described [31]. Briefly, eggs were hatched in deoxygenated, deionized water with powdered Tetramin fish food and larvae were fed Tetramin tablets (Tetra) until pupation. Adult mosquitoes were housed with siblings in BugDorm-1 (Bugdorm) cages and provided constant access to 10% sucrose. Adult females were blood-fed on mice for stock maintenance and blood-fed on human subjects for generation of the *Gr3^ECFP/ECFP^* and *Gr3*^*ECFP*/+^ mosquitoes used in experiments. Blood-feeding procedures with live mice and humans were approved and monitored by The Rockefeller University Institutional Animal Care and Use Committee and Institutional Review Board, protocols 15772 and LVO-0652, respectively. Human subjects gave their written informed consent to participate.

#### Gr3 mutant strain

*Gr3^ECFP/ECFP^* mutants used in this study carry a broadly expressed ECFP marker inserted into the *Gr3* locus as described [8]. *Gr3*^*ECFP*/+^ heterozygotes were the offspring of *Gr3^ECFP/ECFP^* males and wild-type (Orlando) females.

### METHOD DETAILS

#### Fly magnetic tethering

For experiments in Figure S1, flies were magnetically tethered as described [32]. Female flies were collected 1-4 days after eclosion, anesthetized on a Peltier stage at ~4°C for 15-30 min, and attached by the dorsal part of their prothorax to a steel pin using blue-light activated glue (Bondic, Canada). For 1-4 hr after tethering, flies recovered in a dark, humid chamber while holding small squares of tissue paper.

Flies were then suspended in a vertically-aligned magnetic field in the center of a cylindrical LED display (570 nm, IORodeo) covering 360 in azimuth and 94 in elevation with each pixel subtending ~3.75°. LEDs were controlled using PControl in MATLAB [10]. In each experimental bout, flies were exposed to up to six types of visual stimuli composed of LEDs either off (“dark”) or maximally on (luminance 70. cd m^-2^, “bright”), based on previous work [12]: black horizontally centered rectangles 94° tall x 15° wide (“long stripe”), 37×15° (“medium stripe”), and 15×15° (“square”); a uniform bright field (“blank”); a square wave grating composed of 24 alternating bright and dark 15° stripes; and a randomly composed pattern of shuffled dark and bright pixels (“contour”), not analyzed here. The long stripe, medium stripe, square, blank, and contour were randomly presented in 15 s trials, with every 10-16 trials interspersed by a moving 15 s square wave grating stimulus. Each trial presented the shape at a random position on the arena, moving in a sinusoid of peak-to-peak amplitude 60° and frequency randomly chosen from 0, 0.1, 0.2, 0.5, 1.0, 2.0, 4.0, 6.0, or 8.0 Hz. Flies that failed to follow the direction of wide-field motion or failed to sustain flight were discarded.

Flies were lit from below with IR LEDs (850 nm, DigiKey) and video was captured with an infrared-sensitive camera (AVT-GE680) triggered externally, recording frames (320×240 pixels) at 200 Hz. Fly body orientation was extracted from camera images in real time using FView and FlyTrax [33]. These orientation voltages, alongside the camera frame triggers and information about visual stimuli presented, were digitized at 1 kHz using a Digidata 1440a (Molecular Devices).

Data were processed using custom Python software that extracted the offset of each trial by subtracting the center of stimulus position from fly orientation. “Fixation” was calculated by measuring the percentage of time within 3-15 s of stimulus onset spent with an offset between −45° and 45°, while “antifixation” was calculated by measuring the percentage of time within 3-15 s of stimulus onset spent with an offset greater than −135° or 135°. Scores of all trials of each shape were averaged to obtain one score per shape per fly. Because not all flies experienced all four analyzed shapes, we treated fixation scores of the shapes as independent groups for statistical purposes. Heat maps of offsets from trials, separated by shape, show orientation towards the long stripe and away from the spot. Each sector represents 15° x 1 s and are normalized by column.

#### Mosquito magnetic tethering

For experiments in Figure 1 and Figure S1, female mosquitoes were collected 4-15 days after eclosion and fasted in the presence of a water source comprising a 60 mL glass bottle (Fisherbrand™ Clear Boston Round Bottles Without Cap Fisher Sci Cat# 02-911-944) filled with deionized water and plugged with a water-soaked cotton wick (Richmond Dental Braided Rolls ½” x 6”, Catalogue # 201205) for 18-25 hr. Mosquitoes were anesthetized on ice (4°C) for 5-35 min and were attached by the dorsal part of their prothorax to a steel pin using blue-light activated glue (Bondic, Canada). For 1-4 hr after tethering, mosquitoes recovered in a dark, humid chamber while their legs lightly contacted mesh.

Mosquitoes were then suspended in a vertically-aligned magnetic field in the center of a cylindrical LED display (525 nm, IORodeo) covering 360 in azimuth and 94 in elevation with each pixel subtending ~3.75°. LEDs were controlled using PControl in MATLAB [10]. We generated a textured background consisting of pixels randomly assigned to luminances of 10. or 40. cd m^-2^, and we used the same background in the same position for all trials. The long stripe, medium stripe, spot, and blank as used in the fly trials were pseudorandomly presented in 10-15 s trials, with every 8 trials interspersed by 10-15 s of moving the background alone. The long stripe, medium stripe, spot, and blank were superimposed on the background at a random position, moving in a sinusoid of peak-to-peak amplitude 60° and frequency randomly chosen from 0, 0.1, 0.5, and 1 Hz. Mosquitoes that failed to follow the direction of wide-field motion or failed to sustain flight were discarded.

To obtain body orientation, mosquitoes were lit from below with IR LEDs (850 nm, DigiKey) were captured with an infrared-sensitive camera (Point Grey FL3-GE-03S1M-C) triggered externally, recording frames (648 x 488 pixels) at 100 Hz. Mosquito body orientation was extracted from camera images in real time using FView and FlyTrax [33]. Camera frame triggers and information regarding the visual stimuli presented were digitized at 1 kHz using a DAQ (Measurement Computing USB-204) and DAQFlex software (Measurement Computing).

Data were processed using custom Python software that synchronized orientation data and DAQ data using computer timestamps, then processed the same way as fly magnetic tether data. Because all mosquitoes experienced all four analyzed shapes, we treated fixation scores of the shapes as dependent groups for statistical purposes.

#### Glytube blood-meal feeding

For experiments in Figure 1, females were fed sheep blood or saline in groups of 20–50 using Glytube membrane feeders as described [34]. The protein-free saline meal contained 110 mM NaCl, 20 mM NaHCO3, and 1 mM ATP. Glytubes were placed on top of mesh on the mosquito cage, and females were allowed to feed through the mesh for 15 min. Fed females were scored by eye for complete engorgement. Partially-fed females were treated as non-fed and discarded. Blood-fed and saline-fed females were returned to standard rearing conditions. For the “48 hr post-meal” condition, mosquitoes were fasted in the presence of a water source 20-26 hr before testing and then tested 44-52 hr after Glytube feeding. For the “96 hr post-meal” condition, mosquitoes were fasted without a water source 23-49 hr before testing and tested 94-103 hr after Glytube feeding. Water was not provided because this would stimulate females to lay eggs, and the experimental design required females to be gravid at the time of testing.

#### Heat-seeking assay

Experiments in Figure 2, Figure S2, and Figure 3 were performed as previously described [4]. Briefly, the assay apparatus is a 30 x 30 x 30 cm Plexiglass box with a 6 x 9 cm Peltier element (Tellurex) on one vertical wall. To affix a visual stimulus to the Peltier, a 2 cm black dot representing 5.42% of the Peltier area was printed onto a piece of standard white letter size printer paper (extra bright, Navigator; Office Depot/Office Max), which was cut to 15 x 17 cm and held taut over the Peltier by a magnetic frame. For blank control trials in Figure 2 and Figure S2, the paper was turned over to show the unprinted side. This was done to control for the effect of the Xerox Phaser solid printer ink used to generate the dot, which we speculated might affect heat transfer at that position on the paper. Although the dot was faintly visible to the human eye, mosquitoes showed little or no preference for that position on the paper, and there were no detectable differences in the thermal image of the Peltier when the dot was in place (Fig. 2C). All stimulus periods lasted 3 min, followed by 9 min of ambient temperature. CO_2_ pulses (20 s) accompanied all stimulus period onsets. A second identical control Peltier element was situated on the wall opposite to the stimulus Peltier and was set to ambient temperature during all experiments.

Female mosquitoes were separated 8-12 days after eclosion from mixed-sex cages and sorted at 4°C into groups of 45-50. They were kept in custom canisters and sugar-starved in the presence of a water source 13-24 hr before testing. Experiments in Figure 3 were double-blinded to genotype. For each trial, one group was introduced into the enclosure, and only mosquitoes directly on the Peltier area (Fig. 2B) were scored. Due to low contrast between the dark mosquitoes and the black dot, the number of mosquitoes on the dot were scored by hand from videos. Heat maps are smoothed 2D histograms of mean mosquito occupancy during seconds 90–180 of stimulus periods, sampled at 1 Hz and binned into 12 × 16 image sectors.

We quantified positions of landings and takeoffs (Fig. 3E-I) by superimposing two consecutive frames and manually looking for differences. A still mosquito would appear as a completely overlapped image, whereas a walking mosquito would appear as two adjacent images, defined here as two mosquito images that either shared the same orientation and were less than one mosquito body length apart or had orientations with a difference of less than 90° and shared a center of rotation. A mosquito that was present in the first frame but not in an adjacent image in the second frame was counted as a “takeoff”. Conversely, a mosquito present in the second frame but not the first was counting as a “landing” (Fig. 3E). Because we sampled at 1 Hz, it is possible that a seemingly walking or still mosquito represents a mosquito that took off in the first frame and then another mosquito that landed within 1 s in the same or an adjacent location. But we expect that such an event would be exceedingly rare. Thus, this manual quantification represents a conservative estimate of landings and takeoffs. Frames were scored blind to genotype.

Because we did not track individual mosquito identity, we could not track duration of individual dwelling events. However, given the times of landing and takeoff events, we reasoned that we could deduce a mean population dwell time as follows (Fig. 3F). Let A = landing time, B = takeoff time. B – A thus represents dwell time.

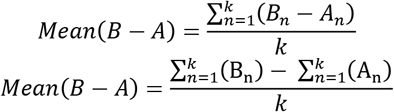

To make this calculation, we assigned all mosquitoes present at the start of the heat epoch a landing time of 0 s and all mosquitoes present at the end of the heat epoch a takeoff time of 180 s.

### QUANTIFICATION AND STATISTICAL ANALYSIS

All statistical analysis was performed using the Python package scipy.stats. Summary data are shown as median with individual data points, and timeseries data are shown as median with median absolute deviation as range. Kruskal-Wallis test with post hoc Mann-Whitney test was used to compare more than 2 independent groups, and Friedman’s test for repeated measures with post hoc Wilcoxon matched-pairs signed-rank test was used to compare more than 2 dependent groups. Mann-Whitney U test was used to compare 2 independent groups. Post hoc tests included Bonferroni correction when multiple comparisons were made. Details of statistical methods are reported in the figure legends.

### DATA AND SOFTWARE AVAILABILITY

Data file containing all raw magnotether data, all processed data, and all code used to process data in this paper are available for download at https://github.com/VosshallLab/LiuVosshall2019. Due to file size limitations, raw data from the heat-seeking assay are available upon request.

